# Mapping the druggable targets displayed by human colonic enteroendocrine cells

**DOI:** 10.1101/2024.10.29.620704

**Authors:** Yuxian Lei, Bettina Bohl, Leah Meyer, Margot Jacobs, Naila Haq, Xiaoping Yang, Bu’ Hussain Hayee, Kevin G Murphy, Parastoo Hashemi, Gavin A Bewick

**Affiliations:** Department of Diabetes, School of Cardiovascular and Metabolic Medicine & Sciences, King’s College London, Guy’s Campus, London SE1 1UL, UK; Diabetes Endocrinology and Obesity Clinical Academic Partnership Kings Health Partners, London, UK; Department of Bioengineering, Imperial College London, South Kensington Campus, London SW7 2AZ, UK; Department of Metabolism, Digestion and Reproduction, Faculty of Medicine, Imperial College London, London W12 0NN, UK; Proteomics Core Facility, James Black Centre, King’s College London (KCL), SE5 9NU, London, UK; Gastroenterology, King’s College Hospital, London, UK

## Abstract

Enteroendocrine cells (EECs) are specialized intestinal hormone-secreting cells that play critical roles in metabolic homeostasis, digestion, and gut-brain communication. They detect diverse stimuli including endocrine, immune, neuronal, microbial, and dietary signals, through a complex array of receptors, ion channels, and transporters, to modulate the release of over 20 hormones. These molecular sensors serve as potential drug targets to modulate hormone secretion, but until recently, catalogues of such targets in human colonic EECs have not been produced.

To address this gap, we performed bulk and single-cell RNA sequencing on fluorescently labelled EECs isolated from human colonic organoids, identifying and cataloguing potential druggable targets. This catalogue includes receptors, orphan GPCRs, transporters, and hormones not previously reported in human colonic EECs. Comparison with murine EECs highlighted interspecies similarities and differences, key data to facilitate the design and optimise the predictive accuracy of pre-clinical models. We also functionally validated two receptors not previously identified in human EECs: IL-13Rα1, was expressed in both peptide-producing EECs and serotonin producing Enterochromaffin cells (ECs), and its ligand IL-13 stimulated the secretion of glucagon-like peptide-1 (GLP-1) and serotonin measured in real-time, and GPR173, which was selectively expressed in ECs and, when activated by its agonist Phoenixin-20, also promoted serotonin release.

These analyses provide a valuable resource for therapeutic interventions aimed at modulating gut hormone secretion, with potential applications in treating gastrointestinal, metabolic, and other related disorders.

## Introduction

Enteroendocrine cells (EECs) are vital sensors and signal transducers within the gastrointestinal epithelium. Although they comprise only ∼1% of epithelial cells, EECs form an extensive hormone-producing network capable of converting luminal, neural, and immune signals into hormonal responses ^1^. EECs have traditionally been classified by the predominant hormone they secrete, e.g., L-cells produce glucagon like peptide-1 (GLP-1), Glucagon like peptide-2 (GLP-2), and Peptide YY (PYY); K-cells secrete Gastric inhibitory polypeptide (GIP); and enterochromaffin cells (ECs) release serotonin (5-HT). Recent advances using fluorescent reporter mice revealed that EEC profiles are shaped by their differentiation trajectory, gut location, and positioning along Wnt and BMP gradients ^2,3,4^. EECs respond to various luminal stimuli including nutrients, pH, and microbial metabolites, releasing over 20 peptide hormones to control that regulate digestion, metabolism, microbial-host cross talk, gut-brain communication and immune defence. Consequently, EEC dysregulation has been implicated in obesity and diabetes, gastrointestinal diseases, inflammation, autoimmune conditions, and certain cancers ^5,6^.

Given their diverse roles, EECs are valuable therapeutic targets and are thought to be important mediators of the metabolic benefits observed in bariatric surgery. Currently, several long-acting variants of individual or combined gut hormones such as GLP-1 and GIP are in development or approved as therapies for obesity and type 2 diabetes ^7^. However, gaining deeper insights into EEC function and how these cells sense stimuli may uncover new druggable targets, enabling more precise control of therapeutic gut hormone secretion from specific EEC subtypes or gut regions. This could broaden therapeutic approaches and introduce new treatment options, including dietary interventions or small molecule drugs. Dietary strategies have the advantage of being more suitable for prevention or population-level interventions and could be more affordable for healthcare systems with limited funding.

Despite their significance, studying human EECs has been difficult due to their sparse distribution and challenges of isolating and maintaining them in culture. Much of our understanding is based on murine models, where fluorescent labelling techniques have mapped EEC distribution. However, the development of organoid technology and CRISPR-Cas9 gene editing has revolutionized the field. These advances allow us to grow three-dimensional human intestinal organoids and study EECs in unprecedented detail ^8,9^.

We have taken advantage of this technology to map druggable targets expressed in human colonic EECs. We compared this dataset with data from mouse colonic EEC subtypes to probe interspecies similarities and differences and identified novel receptors enriched in human colonic EECs. Among those, we selected IL-13Rα1 and GPR173, and corroborated their effects on hormone secretion ^10^. By providing a map of human colonic EEC cell surface targets we hope to facilitate the identification of novel EEC biology and therapeutic targets.

## Results

### Generation of human colonic CHGA reporter organoids

Chromogranin A (CHGA) is a marker of neuroendocrine cells and marks EECs in the gut epithelium. We generated human CHGA-mNeon expressing colonoids using CRISPR–Cas9-mediated homology-independent transgenesis (CRISPR-HOT), as previously described ^9,11^. The fluorescent tag mNeon was inserted 7 base pairs prior to the stop codon (TGA) of the endogenous *CHGA* gene (**Fig. 1a**). To enrich for both peptide producing EECs and ECs, we combined two previously published differentiation protocols. This involved the use of differentiation medium which consisted of insulin-like growth factor 1 (IGF-1) and fibroblast growth factor 2 (FGF-2), with the removal of epidermal growth factor (EGF) ^12,13^, and a 48-hour pulse treatment with a Notch inhibitor (iNotch), MEK inhibitor (iMEK), and ISX-9 ^14^ (**Fig. 1b**). This combination significantly increased EEC and EC differentiation.

**Fig. 1.**
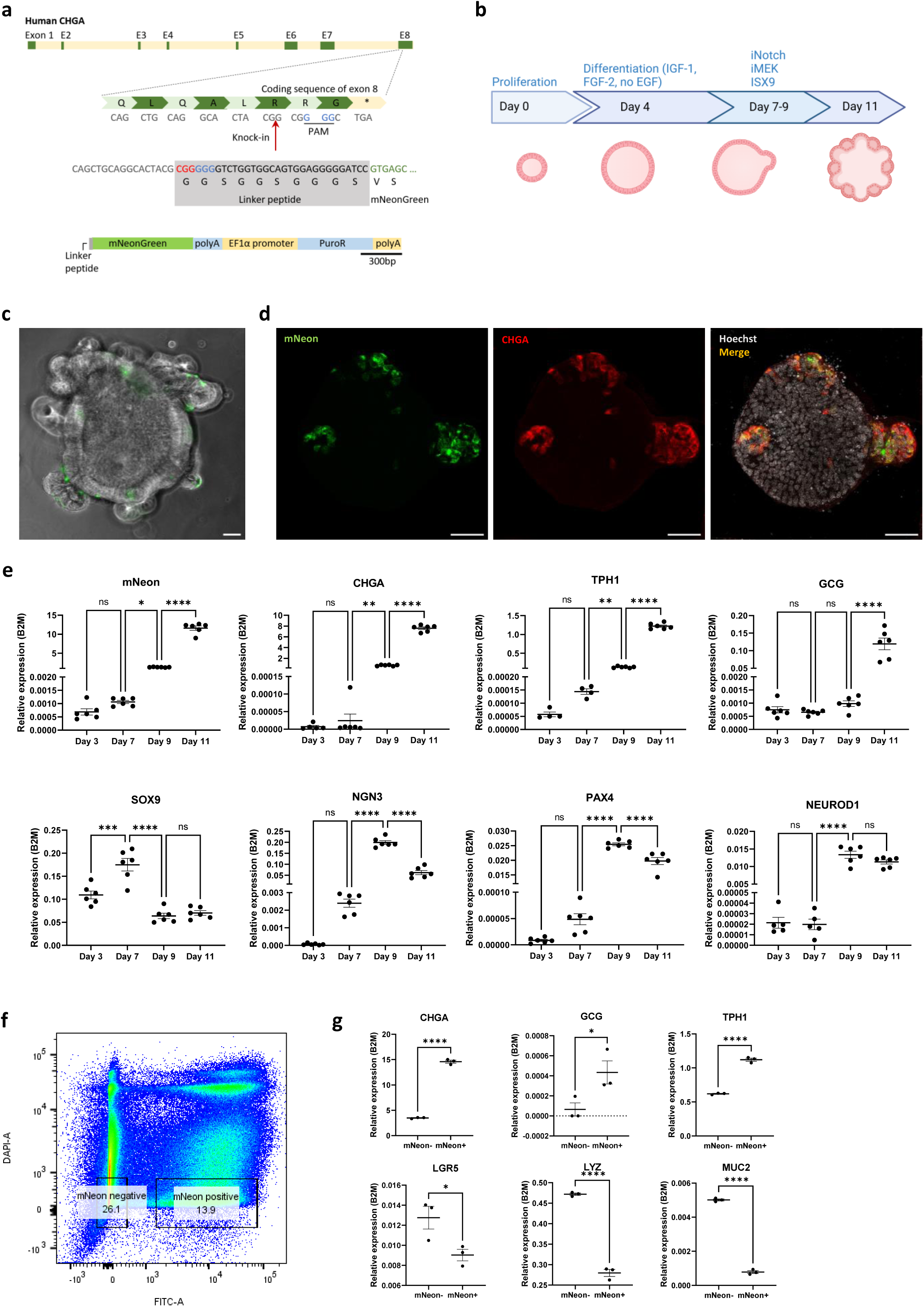
Generation of human CHGA-mNeon reporter colonoids. **a,** Schematic showing mNeon insertion into the CHGA gene by CRISPR–Cas9-mediated homology-independent organoid transgenesis (CRISPR-HOT). **b,** Schematic demonstrating differentiation protocol of human colonoids to enrich enteroendocrine cells (EECs). **c,** Live image of CHGA-mNeon colonoids showing the mNeon green fluorescence reporter expression. Scale bar, 50 µm. **d,** Confocal images of immunofluorescent staining of CHGA-mNeon human colonoids for CHGA (red), which colocalized with mNeon green fluorescence. Scale bar, 50 µm. **e,** qPCR analysis of key EEC developmental markers in the time course differentiation of CHGA-mNeon human colonoids. Data are represented as mean ± SEM. *p < 0.05, **p < 0.01, ****p < 0.0001 by one-way ANOVA tests. **f,** Representative fluorescence-activated cell sort (FACS) plot of 500,000 events. mNeon-positive and -negative cells were sorted based on mNeon fluorescence, with DAPI-negative gating. **g,** qPCR analysis of mNeon^-ve^ and mNeon^+ve^ sorted cells. Data are represented as mean ± SEM. *p < 0.05, ****p < 0.0001 by unpaired t tests.

Differentiated CHGA-mNeon colonoids displayed scattered green-fluorescent cells throughout their structure (**Fig. 1c**). Colocalization of CHGA and mNeon was confirmed by immunostaining (**Fig. 1d**), indicating successful tagging of CHGA-expressing cells. We evaluated our differentiation method using time-course qPCR of whole colonoids (**Fig. 1e**). The stem cell marker *SOX9* and the endocrine progenitor markers *NGN3* and *PAX4* were upregulated early in the differentiation process, with the endocrine progenitor markers showing a slight delay relative to *SOX9*. These markers were downregulated later in the differentiation time course. Conversely, markers of mature endocrine cells (*CHGA*, *NEUROD1*, tryptophan hydroxylase 1 (*TPH1*), preproglucagon (*GCG*)) increased over time, peaking on day 11. As expected, *mNeon* expression mirrored *CHGA* expression. Together the data suggested our protocol successfully enriched human colonoids with EECs and ECs.

CHGA-mNeon positive cells were purified from differentiated colonoids using fluorescence-activated cell sorting (FACS), yielding two populations: fluorescent mNeon^+ve^ and non-fluorescent mNeon^-ve^ cells (**Fig. 1f**). The mNeon^+ve^ population was enriched for EEC markers (*CHGA*, *GCG*) and the EC marker *TPH1* (**Fig. 1g**). Conversely, mNeon^-ve^ cells were enriched for Paneth cell (*LYZ*), intestinal stem cell (*LGR5*), and goblet cell (*MUC2*) markers. Proteomic analysis confirmed that mNeon^+ve^ and mNeon^-ve^ cells were significantly separated at the level of protein expression (**Fig. S1a**). mNeon^+ve^ cells produced CHGA, TPH1 and GCG, whereas mNeon^-ve^ cells produced proteins such as LYZ and the goblet cell marker TFF3 (**Fig. S1b**). These expression patterns demonstrate that mNeon^+ve^ cells are human colonic EECs and ECs.

### Mapping the transcriptome of human colonic EEC and ECs

To characterize the transcriptome of mNeon^+ve^ cells, we performed bulk RNA-sequencing (RNA-seq). Principal component analysis (PCA) confirmed clear transcriptional separation between mNeon^+ve^ and mNeon^-ve^ populations (**Fig. 2a**). The mNeon^+ve^ population exhibited EEC marker genes like *CHGA*, *CHGB*, and *SCG5* (**Fig. 2b**), and transcription factors (TFs) associated with EEC development including *FEV*, *INSM*, *NKX2-2*, *NKX2-1* (**Fig. S1c**). As expected, the enterocyte marker *SLC26A3*, the Paneth cell markers *CA4*, *SPIB*, and the goblet cell markers *CLCA1*, *MUC5AC*, *SPDEF*, *SPINK4*, and *TFF3* were enriched in mNeon^-ve^ cells. Hormone-encoding genes, including *GCG*, which produces the incretin GLP-1, and the anorexigenic peptide YY (*PYY*), were enriched in mNeon^+ve^ cells. These are characteristic products of peptide producing EECs. Additionally, *TPH1*, the rate-limiting enzyme in the synthesis of serotonin and a marker of ECs, was also enriched, along with transcription factors associated with EC differentiation such as *PAX4* ^15^ and *RUNX1T1* ^2^. Other classical gut hormones, secretin (*SCT*) and neurotensin (*NTS*), were expressed at lower levels.

**Fig. 2.**
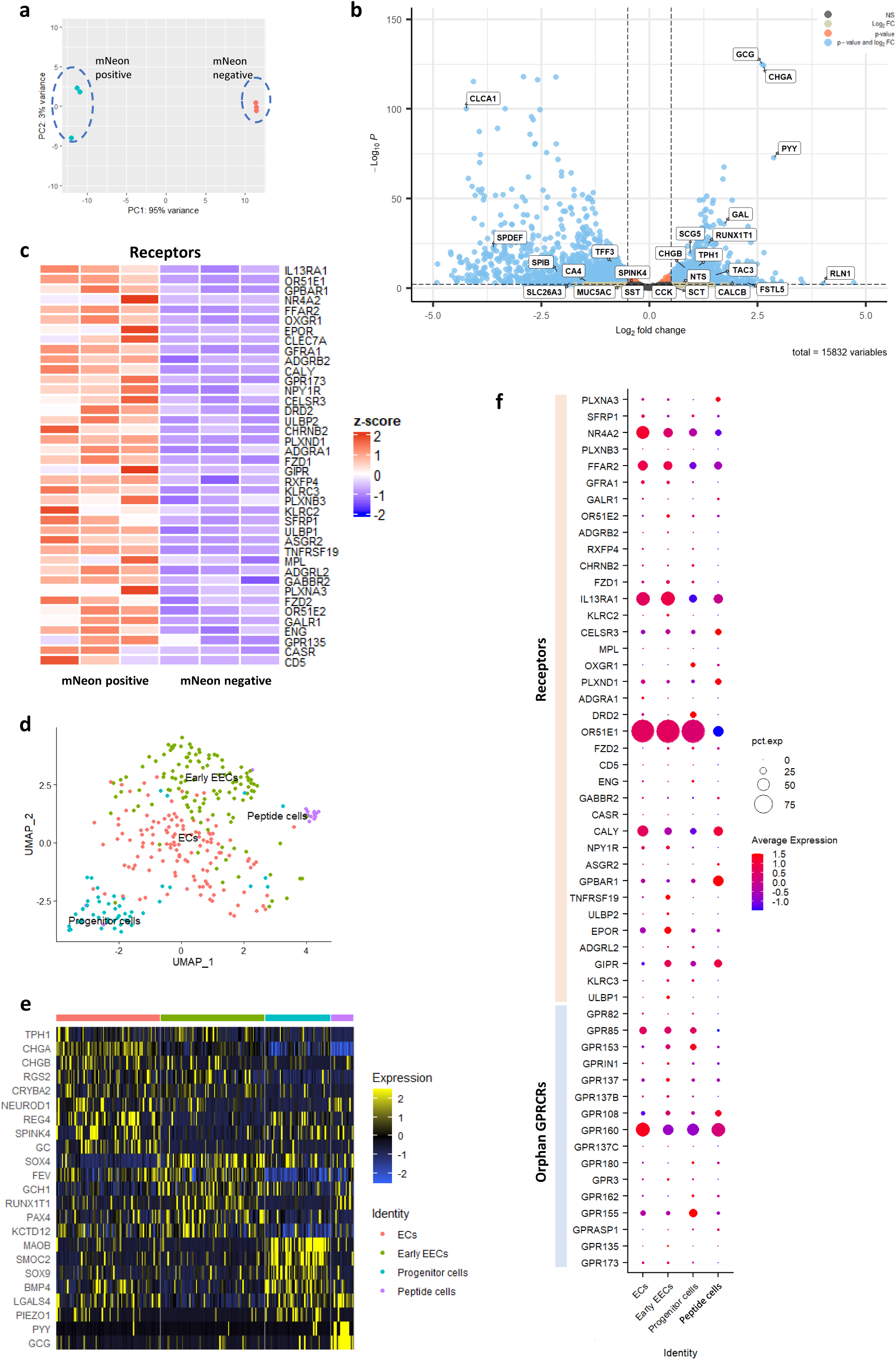
Transcriptomic analysis of CHGA-mNeon cells on bulk and single-cell scales. **a,** Principal component analysis (PCA) plot comparing mNeon positive and negative cell populations by bulk RNA-sequencing (n=3). **b,** Volcano plot showing differential expression of selected genes. **c,** Heatmap showing differentially expressed receptor genes with the highest fold change and a p value < 0.01. **d,** UMAP showing segregation and annotation of CHGA-mNeon cells by single-cell RNA-sequencing. **e,** Heatmap showing relative expression of selected key developmental markers for the EECs across different cell clusters. **f,** Dot plot of genes coding receptors and orphan GPCRs, identified across different human colonic EEC clusters. Size of the circles represents percentage of cells expressing the gene and color of the circles represents average expression of indicated genes partitioned by clusters.

Beyond the well-known gut hormones, our analysis of genes predicted to produce secreted products identified several additional neuropeptides and hormones not previously reported in human colonic peptide EECs or ECs (**Fig. S1c**). These included galanin (*GAL*), a potent inhibitor of gut hormone secretion (including GLP-1, PYY, and NTS) and a known regulator of energy homeostasis and intestinal motility ^16,17^. Additionally, we found previously unreported expression of relaxin 1 (*RLN1*), which modulates the reproductive system and gut motility ^18^; calcitonin gene-related peptide (*CALCB*), a neuropeptide with functions in the gut including the regulation of mucosal blood flow, epithelial homeostasis, exocrine and endocrine secretory processes, motor activity, and nociception ^19^; and neurokinin B (*TAC3*), a member of the tachykinin family of neuropeptides involved in reproduction, gastric acid secretion, GI motor and sensory control, and inflammation ^20,21^. Interestingly, our analysis also revealed the presence of follistatin-like 5 (*FSTL5*), a secretory glycoprotein previously implicated in the olfactory system ^22^. While its role in the gut is unclear, it has been shown to inhibit proliferation in liver hepatocellular carcinoma ^23^.

Among the top differentially expressed receptors (**Fig. 2c**) were the bile acid receptor (*GPBAR1*) and free fatty acid receptor 2 (*FFAR2*), both implicated in the regulation of postprandial nutrient sensing and gut hormone secretion ^24^. These receptors have been reported to be expressed by EECs and/or ECs. Several hormone receptors were also enriched, including the gastric inhibitory polypeptide receptor (*GIPR*). Its ligand, GIP, is an incretin secreted from K-cells in the small intestine. Our data corroborated the presence of the neuropeptide Y receptor type 1 (*NPY1R*), which has previously been identified on colonic goblet cells, endocrine cells, and enterocytes ^25^. NPY1R mediates some effects of the NPY-like family of peptides on gut functions, including secretion of GLP-1, inflammation, barrier function, and absorption ^26–29^. Additionally, the galanin receptor 1 (*GALR1*) and the relaxin family peptide (*RXFP4*) were also present. RXFP4 and its ligand, insulin-like peptide 5 (INSL-5), are thought to play a role in the regulation of food intake and gut motility. Recent data revealed co-localization of RXFP4 with serotonin in the lower gut of mice and single cell RNA-sequencing has suggested its presence in a subset of human colonic ECs ^30^.

EECs and ECs are strategically positioned to receive information from various sources, including enteric and vagal neurons. They can respond to classical neurotransmitters released by these neurons, and to dietary and microbial signals that in some instances act as ligands for neurotransmitter receptors. Our data demonstrate the presence of the γ-aminobutyric acid (GABA) receptor (*GABBR2*), the cholinergic receptor (*CHRNB2*) and the dopamine receptor D2 (*DRD2*) in mNeon^+ve^ cells (**Fig. 2c**), extending previous findings in mice to humans ^31^. Furthermore, we confirmed the presence of the olfactory receptors (*OR51E1*, *OR51E2*) in human colonic endocrine cells ^8^.

EECs and ECs play a crucial role in orchestrating gut epithelial immunity and inflammation. They possess multiple receptors which allow them to respond to immune mediators and can directly secrete cytokines and other modulators. For example, we found the receptors: interleukin-13 (IL-13) receptor *IL13RA1*, *ULBP1*, *CLEC7A*, *KLRC3*, *TNFRSF19*, and *CD5* (**Fig. 2c**), and cytokines including *CCL5*, *CCL15*, *CSF1*, *IL17*, *IL11*, *IL23* (**Fig. S1c**), and various tumour necrosis family members, enriched in mNeon^+ve^ cells. The presence of *IL13RA1* and *IL17* corroborates the known role of IL-13 in controlling ECs in mice ^32^ and the previously reported production of IL-17C by EECs in humans ^33^.

We also identified differentially expressed transporters consistent with the chemosensing role of EECs and ECs (**Fig. S1c**), including *SLC22A17* (iron transporter), *SLC38A11* (amino acid transporter), *SLC17A9* (ATP uniporter), *SLC26A7* (anion exchange transporter), and *SLC8A1,* a sodium/calcium exchanger. As expected, we identified numerous enriched ion channels associated with excitable cells, such as potassium channels (*KCTD12*, *KCNJ3*, *KCNJ6*, *KCHN2*), calcium channels (*CACNA2D1*, *CACNA1A*, *CACNA1C*), and sodium channels (*SCNA3*, *SCN3B*, *SCNA2*).

To complete our survey of druggable targets, we identified the top differentially expressed orphan G protein-coupled receptors (**Fig. S1c**), which to our knowledge have not been previously reported in either mouse or human colonic EECs or ECs. Interestingly, within this list of novel targets, GPR173 has recently been deorphanized and is proposed to be the receptor for Phoenixin (PNX), a recently discovered neuropeptide involved in the regulation of food intake, energy homeostasis, stress and inflammation ^34^.

### Single-cell transcriptomic profiling of human colonic EECs and their receptor expression

Enriching human EECs from primary tissues for single-cell profiling has been challenging due to reliance on antibodies and selection strategies that can lack specificity. To expand on our bulk sequencing data, we performed single cell RNA-sequencing on our CHGA reporter organoids, to profile mNeon^+ve^ cells, focusing on the expression of druggable cell surface targets within specific cell populations. Data was generated and processed by sorting and robot-assisted transcriptome sequencing (SORT-seq ^35^) and analysed using Seurat ^36^. Using shared nearest neighbour clustering based on highly differentially expressed genes, we constructed a map of CHGA-mNeon^+ve^ cells. The map contained four populations of colonic EECs: progenitor cells, early EECs, peptide EECs and ECs (**Fig. 2d**).

These findings align with the two major branches of differentiated endocrine cells: TPH1-positive ECs and hormone expressing (e.g., GLP-1 positive) EECs, as previously documented ^25^. Progenitors were identified by the expression of the stem cell markers *SMOC2*, *SOX9* and *BMP4*, and *LGALS4*, a marker of the early NGN3-to-EC transition ^37^ (**Fig. 2e**). Early EECs were distinguished by the expression of *PAX4*, *SOX4*, *GCH1* and *RUNX1T1* ^8,37^; this population likely harbours cells with the potential to become ECs as well as hormone producing EECs but with an EC bias due to the presence of *FEV* and *PAX4*. Interestingly, *PAX4* has recently been identified as a critical regulator in an endocrine transcription factor network controlling EC differentiation ^15^ and is one of the major targets of ISX-9, a component of our differentiation protocol. The late EC population was identified by enrichment for *CHGA*, *CHGB* and *TPH1* and genes such as *REG4, GC, RGS2,* and *CRYBA2* which have previously been observed at the later stages of differentiation from progenitor to mature ECs in humans ^37^. The EEC peptidergic cluster was identified based on the expression of classical markers, including *GCG*, *PYY* and *NeuroD1*.

Single-cell clustering identified the most highly expressed receptors and orphan GPCRs enriched in colonic EECs, with variable expression across cell subtypes (**Fig. 2f**). Receptors most highly expressed in ECs included nuclear receptor subfamily 4 group A member 2 (*NR4A2*), a regulator of inflammation in the gastrointestinal tract ^38^, the olfactory receptor *OR51E1*, and the Phoenixin receptor *GPR173*. In contrast, receptors such as cadherin EGF LAG seven-pass G-type receptor 3 (*CELSR3*), a key gene for planar cell polarity and the guidance of enteric neuronal projections ^39^, the bile acid receptor (*GPBAR1*), gastric inhibitory polypeptide receptor (*GIPR*), galanin receptor 1 (*GALR1*), and the orphan receptor *GPR108* were primarily enriched in peptide-producing EECs. Receptors with broader expression included *GPR160*, an orphan receptor potentially activated by CART (cocaine and amphetamine-regulated transcript) peptide, a neuropeptide and hormone involved in appetite regulation ^40^, gastrointestinal inflammation, and gut hormone secretion from the proximal bowel (GLP-1 and GIP) ^41^, as well as *IL13RA1* (interleukin-13 receptor subunit alpha-1). *GPR155* showed the highest expression in progenitor cells and has been linked to cancer initiation and progression, particularly as a biomarker for hematogenous metastasis in gastrointestinal cancers ^42^. However, its role in regulating intestinal stem cells (ISCs) or progenitor EEC proliferation and differentiation remains unclear. Additionally, 2-oxoglutarate receptor 1 (*OXGR1*) was enriched in progenitor cells. Dietary and gut microbiota-derived 2-oxoglutarate is utilized by the epithelium for protein synthesis and oxidative metabolism and has been shown to restore intestinal barrier function by promoting stem cell activity in mice ^43^.

### An integrated transcriptomic database of mouse and human colonic EECs

To compare interspecies differences and similarities between humans and mice, we conducted transcriptomic analysis of mouse colonic EECs. We used mouse TPH1 and PYY reporter cells, which are analogous to human colonic ECs and peptide hormone producing L-cells, respectively. TPH1^+ve^ cells were isolated from colonoids derived from Tph1-P2A-iCreERT2 mice ^44^ (**Fig. 3a**). Principal component analysis (PCA) revealed two distinct CFP^+ve^ and CFP^-ve^ cell populations (**Fig. 3d**). CFP^+ve^ cells were enriched for the EC marker *Tph1*, whilst CFP^-ve^ cells were enriched with the Paneth cell marker *Lyz* (**Fig. 3b, c**). As expected, CFP^+ve^ cells were also enriched with EEC lineage markers such as *Neurog3* and *Nkx2-2* (**Fig. 3e**). These cells also showed enrichment for transcription factors driving EC differentiation, including *Pax4* and *Runx1t1*, along with TFs involved in serotonin biosynthesis, such as *Lmx1a* ^45^ and *Ascl1* ^46^ (**Fig. S2a**). As expected, the epithelial marker *Slc26a3*, goblet cell differentiation factor *Klf4*, and Paneth cell marker *Mmp7* were enriched in CFP^-ve^ cells (**Fig. 3e**).

**Fig. 3.**
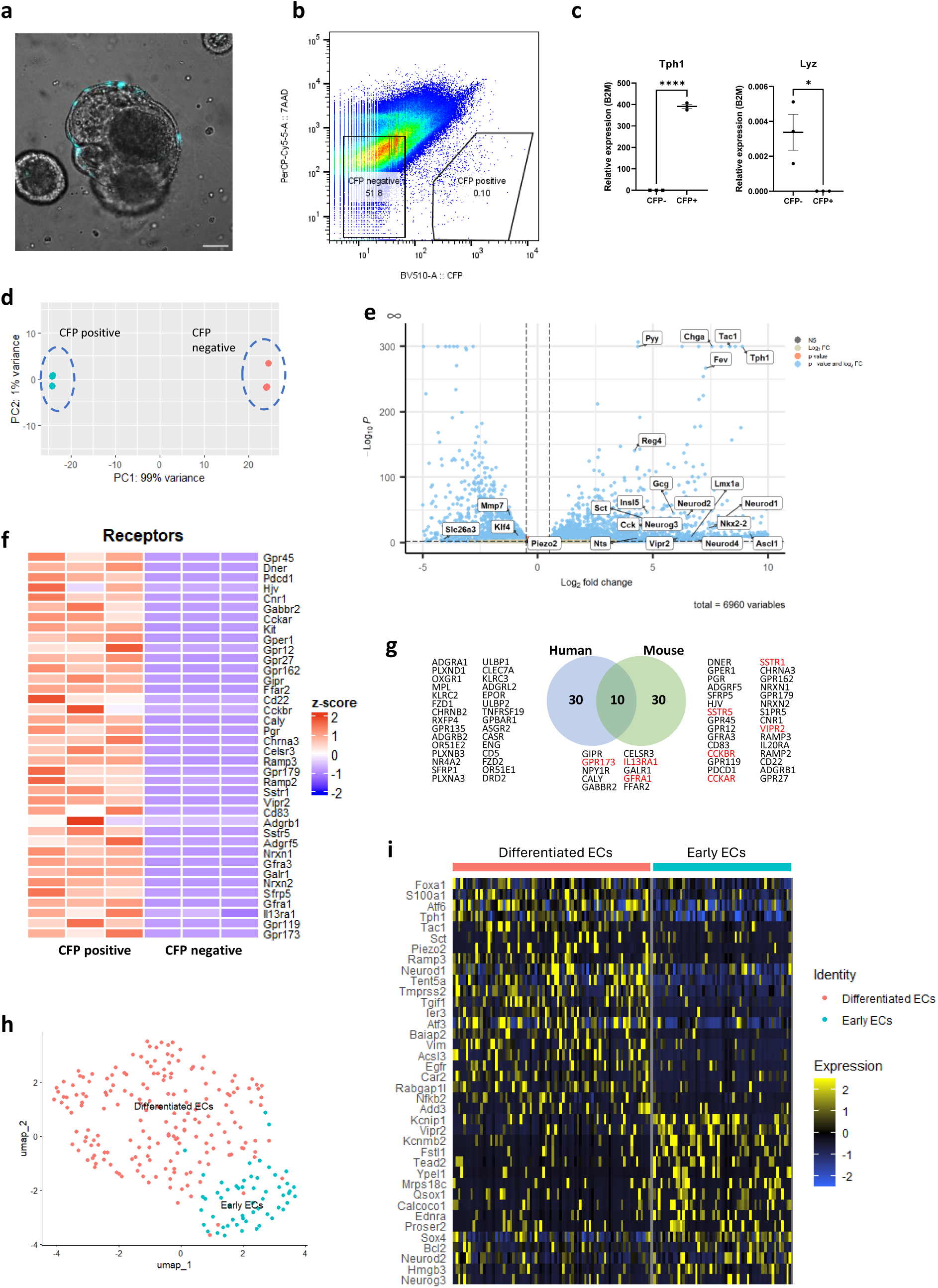
Transcriptomic analysis of mouse colonic Tph1-CFP cells on bulk and single-cell scales. **a,** Live image of mouse TPH1-CFP colonoids showing the CFP fluorescence reporter expression. Scale bar, 50µm. **b,** FACS output plot of 450,000 events, showing the sorting of CFP-positive and -negative cells, with 7AAD-negative gating. **c,** qPCR analysis of CFP^-^ and CFP^+^ sorted cells. Data are represented as mean ± SEM. *p < 0.05, **p < 0.01, ****p < 0.0001 by unpaired t tests. **d,** PCA plot of CFP-positive and -negative cell populations by bulk RNA-sequencing (n=3). **e,** Volcano plot showing differential expression of selected genes. **f,** Heatmap showing differentially expressed receptor genes with the highest fold change and a p value < 0.01. **g,** Venn diagram showing the conserved and uniquely expressed top receptor genes between human EECs and mouse ECs. **h,** UMAP showing segregation and annotation of Tph1-CFP cells by single-cell RNA-sequencing. **i,** Heatmap showing relative expression of selected key markers in Tph1-CFP cells across different cell clusters.

Ninety receptors were differentially expressed in mouse CFP^+ve^ ECs compared to CFP^-ve^ cells, including known receptors involved in nutrient, gut hormone, and immune sensing. Among the top 40 differentially expressed receptors (**Fig. 3f**), 10 were conserved between mouse and human (**Fig. 3g**), including receptors such as *FFAR2*, *GABBR2*, and *NPY1R*, as well as several herein newly identified receptors, including *IL13RA1*, *CELSR3*, and *GFRA1*. Glial cell line-derived neurotrophic factor (GDNF), the ligand for *GFRA1*, is crucial for the development of the enteric nervous system ^47^. Additionally, *GFRA1* has been implicated in Hirschsprung’s disease and related enterocolitis in mice ^48^. However, its expression and role in ECs has not been explored. Several hormone receptors were exclusively expressed in the mouse dataset (**Fig. 3g**), including the vasoactive intestinal polypeptide receptor (*Vipr2*), whose ligand VIP regulates nutrient absorption and gut motility ^49^; the somatostatin receptors (*Sstr5*, *Sstr1*), which play versatile roles in the gut, including chemosensing, mucus secretion, and inflammatory responses ^50^; and the CCK receptors (*Cckar*, *Cckbr*), which regulate gut motility and appetite. Selective species expression may indicate divergent hormonal control of ECs between mouse and humans.

Among the orphan GPCRs (**Fig. S2b**), *Gpr173*, *Gpr162*, *Gpr85*, *Gpr146*, *Gpr153*, *Gpr6*, *Gpr3* were conserved between mouse and human. Sphingosine-1-phosphate (S1P) has been identified as a likely ligand for *Gpr6*, *Gpr3* ^51^, and is implicated in intestinal barrier function ^52^. In contrast, *Gpr12*, *Gpr27*, *Gpr45*, *Gpr161* and *Gpr179* were selectively found in the mouse. Of these *Gpr12* ^53^, *Gpr27* ^54^, *Gpr45* ^55^ have been linked to obesity and/or metabolism, whereas *Gpr161* has a role in intestinal immunity ^56^. In the human mNeon^+ve^ dataset, *Gpr63* and *Gpr68* were receptors with postulated ligands. For example, sphingosine-1-phosphate (S1P) and lysophosphatidic acid (LPA) have been suggested as low affinity agonists of *Gpr63* ^57^. *Gpr68*, also known as the proton-sensing ovarian cancer G-protein coupled receptor (OGR1), is a proton-activated GPCR that responds to acidic extracellular pH and in the intestine plays a role in tissue damage and inflammation ^58^.

There were also various receptors involved in gut inflammation and immune responses (**Fig. 3f, g**), including the conserved receptor Il13ra1 and receptors uniquely expressed in mice, such as Pdcd1 (which encodes the programmed cell death protein 1), ^59^, *Il20ra* ^60^, *Cd83* ^61^, *Cd22* ^62^. Additionally, cytokines, such as the conserved Il17d, as well as mouse specific Ccl6, Ccl28, Mif are also implicated in these processes (**Fig. S2a**). Our analysis revealed both conserved and species-specific expression across other functional groups, highlighting the importance of understanding species-dependent expression to facilitate the optimisation of pre-clinical models for the validation of novel targets (**Fig. S2**).

In addition to the bulk transcriptome analysis of TPH1-CFP^+ve^ cells, we also performed single cell RNA-sequencing using these cells and PYY-GFP cells, isolated from the colon of PYY-GFP mice ^63^ (**Fig. 4a**). The CFP^+ve^ ECs clustered into two groups: differentiated ECs and early ECs, mirroring clusters observed in human ECs derived from colonoids (**Fig. 3h**). Differentiated ECs were identified by the expression of genes such as *Foxa1*, *S100a1* and *Atf6*, which were observed during late EC lineage development in mice ^2^ (**Fig. 3i**). Early ECs demonstrated markers like *Neurog3*, *Neurod2* and *Sox4*. As expected, PYY-GFP cells were enriched for *Pyy* and *Gcg* expression and not for *Tph1* or *Lyz* (**Fig. 4a, b**). PYY-GFP cells clustered into peptide EECs, which highly expressed *Chga*, *Gcg*, and *Pyy*, and secretory progenitor cells, which expressed the markers *Atoh1* ^64^, *Spink4*, and *Agr2*, and exhibited features of both goblet and Paneth cells (**Fig. 4c, d**). The top differentially expressed receptors were compared between the TPH1 and PYY clusters (**Fig. 4e**). *Ffar2* was the most promiscuous, being expressed in all clusters whereas *Gpbar1* was restricted to mature peptide EECs. When expression patterns were compared between human and mouse. *Npy1r* exhibited a conserved expression pattern, with predominant expression in ECs, whilst *Ffar2* was broadly expressed across all EEC clusters in both mouse and humans. However, other receptors exhibited species-specific differences. For example, *Gpr108*, an immune modulator, exhibited the highest expression in human peptide EECs but in the mouse was predominantly expressed in secretory progenitor cells and mature ECs. *Il13ra1* also displayed differential expression, being primarily expressed in mouse secretory progenitor cells and early ECs, while in humans it was expressed across all stages of EC differentiation as well as in peptide EECs. Interestingly, the orphan receptor *Gpr173* was undetected in mouse peptide EECs, consistent with its absence in human EECs, suggesting its expression is conserved and restricted to ECs.

**Fig. 4.**
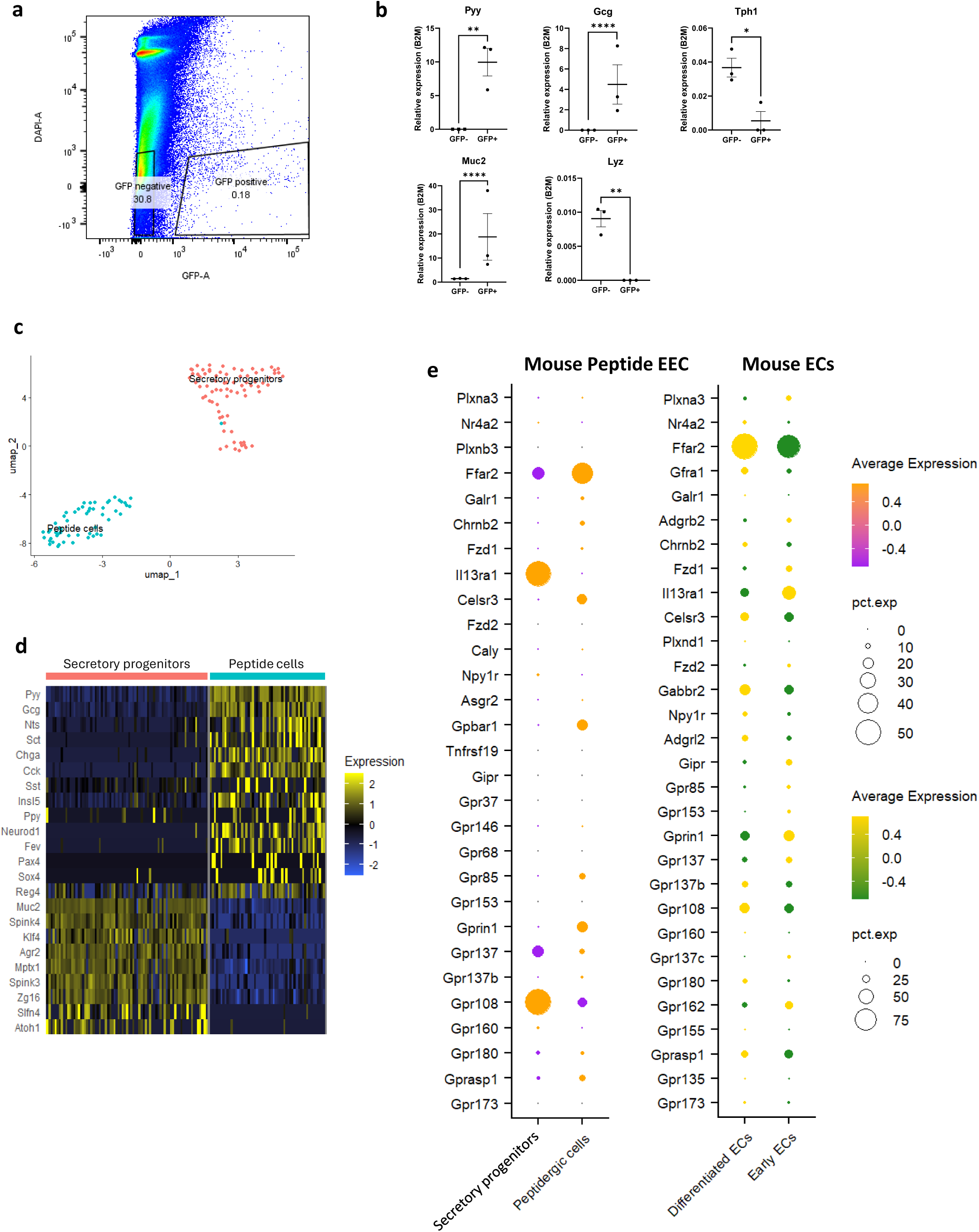
Transcriptomic analysis of mouse colonic Pyy-GFP cells. **a,** FACS output plot of 500,000 events, showing the sorting of PYY-GFP-positive and -negative cells, with DAPI-negative gating. **b,** qPCR analysis of GFP^-^ and GFP^+^ sorted cells. Data are represented as mean ± SEM. *p < 0.05, **p < 0.01, ****p < 0.0001 by unpaired t tests. **c,** UMAP showing segregation and annotation of Pyy-GFP cells by single-cell RNA-sequencing **d,** Heatmap showing relative expression of selected key markers in Pyy-GFP cells across different cell clusters. **e,** Dot plot of genes coding human receptors and orphan GPCRs were aligned across different mouse colonic EEC clusters. Size of the circles represents percentage of cells expressing the gene and color of the circles represents average expression of indicated genes partitioned by clusters.

The complete lists of differentially expressed receptors, transcription factors, and cytokines in mouse ECs and human EECs are provided in Supplementary Tables S1 and S2. Additionally, the comparisons of average gene expression across all clusters of human EECs and mouse ECs are available in Supplementary Tables S3 and S4.

### Are receptors enriched on human EECs functionally significant?

To evaluate whether we could identify receptors with functional significance in both ECs and peptide EECs, we selected IL-13Rα1 and GPR173. IL-13Rα1 has previously been shown to regulate serotonin secretion in mice, but its presence on human L-cells has not previously been reported. Whereas GPR173 has not been reported on ECs in either species. The expression of both receptors was confirmed by qPCR in human CHGA-mNeon^+ve^ cells (**Fig. 5a**). Additionally, co-expression of IL-13Rα1 with GLP-1 and 5-HT was further corroborated in human colon biopsies by immunohistochemistry (**Fig. 5c**). Consistent with its expression in human EECs, *Il13ra1* expression was significantly increased of in both mouse TPH1-CFP^+ve^ cells (ECs) and PYY-GFP^+ve^ cells (peptide EECs) (**Fig. 5d**), while the expression of *Gpr173* was enriched in mouse ECs.

**Fig. 5.**
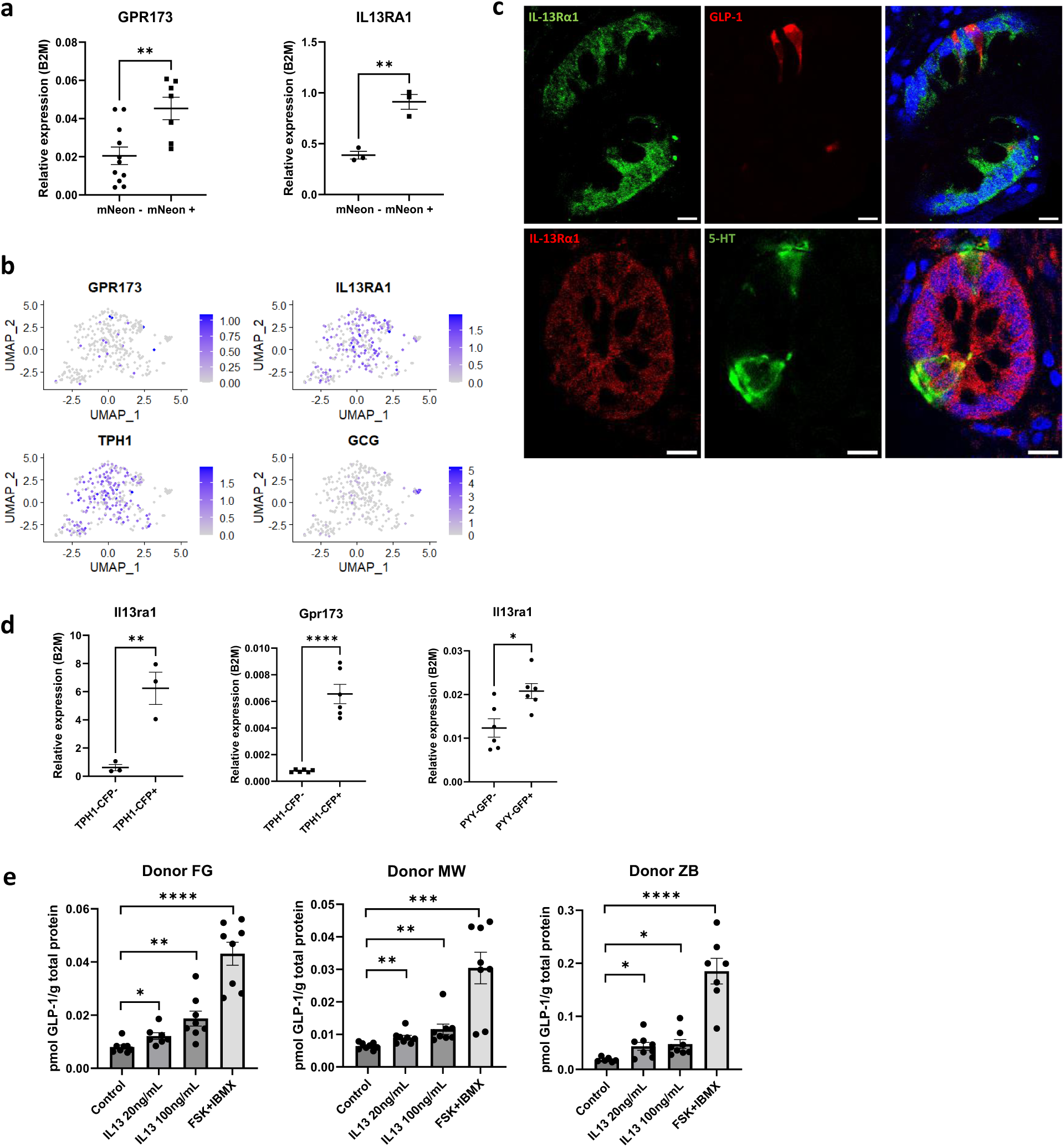
Activation of IL13RA1 induced GLP-1 secretion. **a**, qPCR analysis showing enrichment of receptor genes *GPR173* and *IL13RA1* in human CHGA-mNeon cells. **b,** UMAP displaying co-expression of receptor genes *GPR173* and *IL13RA1* with hormone genes *TPH1* and *GCG* in human colonic EECs on the single-cell level. Bars display color-coded unique transcript expression in logarithmic scale. **c,** Immunohistochemical staining showing colocalization of IL13RA1 with GLP-1 or 5-HT in human colon mucosal biopsy sections. Scale bar 10 µm. **d,** qPCR analysis showing enrichment of *GPR173* in mouse Tph1-CFP cells, *IL13RA1* in both Pyy-GFP and Tph1-CFP cells. **e,** GLP-1 secretions in response to 20 and 100 ng/mL IL-13, and 10 µM forskolin (FSK) plus 10 µM 3-isobutyl-1-methylxanthine (IBMX) were measured in supernatants by ELISA. Values were normalized against the total protein concentrations. Data are represented as mean ± SEM. Unpaired t tests were performed, with *p < 0.05, **p < 0.01, ***p < 0.001, ****p < 0.0001.

The presence of IL-13Rα1 on L-cells suggests that the IL-13/IL-13Rα1 pathway might stimulate the secretion of GLP-1. In support of this, we found that IL-13 significantly increased GLP-1 secretion in colonoids derived from three separate donors (**Fig. 5e**). IL-13 has been shown to stimulate serotonin secretion from BON-1 cells, a human carcinoid cell line that produces serotonin (5-HT) and other neurotransmitters and peptides ^32,65^. However, it remains unknown whether IL-13 can stimulate serotonin secretion from primary human ECs. Accurately measuring serotonin secretion from primary human cells is challenging. Unlike secretion protocols for gut hormones, which rely on sensitive immunoassays, real-time serotonin measurement requires electrochemical techniques that necessitate placing electrodes near the secreting cells. This is impractical without a reporter to mark the cells of interest. Previously, 5-HT release from individual ECs in mice has been indirectly visualized by measuring Ca^2+^ responses in fluorescently labelled ECs in organoids, or by indirectly measuring activation of biosensor cells expressing the serotonin-gated ion channel (5-HT_3_R) ^66^.

Here, we employed Fast Scan Cyclic Voltammetry (FSCV), an electrochemical technique used to simultaneously identify and quantify monoamines, to investigate whether the IL-13/IL-13Rα1 pathway stimulates serotonin secretion from primary human ECs. This method allows for real-time and label-free detection of extracellular 5-HT as a change in current at a carbon fibre microelectrode (CFM) (**Fig. 6a**). Combining with CHGA-mNeon colonoids enabled the measurements to be performed in or near primary ECs. We compared the spontaneous (pre-drug) activity of CHGA-mNeon fluorescent cells to their post-drug stimulation. Representative current-time plots were generated for each organoid (**Fig. 6b**). We quantified the frequency of 5-HT release events and observed significant increases following IL-13 stimulation (**Fig. 6c**). Furthermore, we measured the peak amplitude corresponding to the amount of 5-HT released for each event and found IL-13 treatment increased peak amplitude (**Fig. 6d**). These results suggest that IL-13 increases both the number of 5-HT release events, and the amount of 5-HT released in each event.

**Fig. 6.**
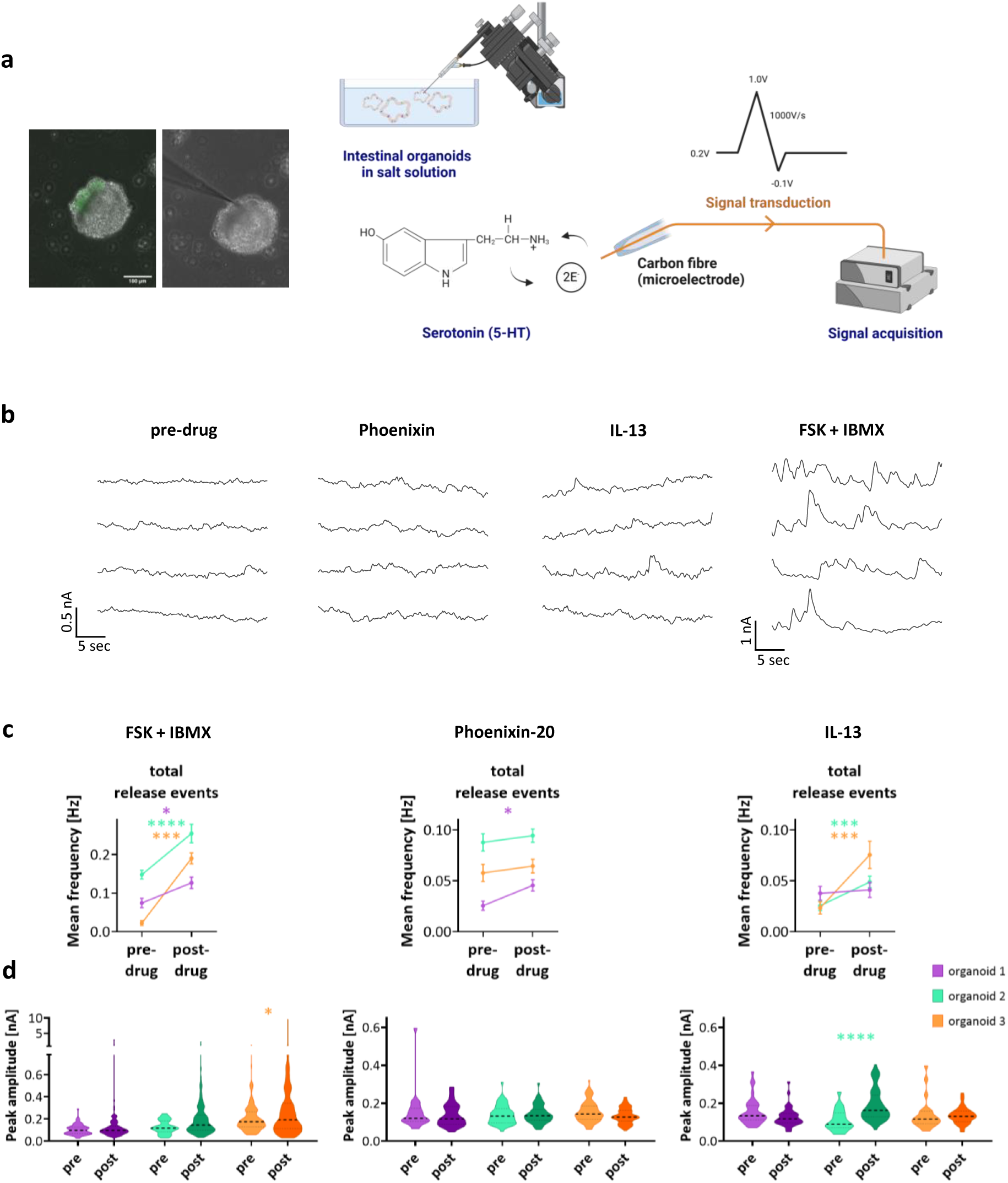
Activation of IL13RA1 and GPR173 induced serotonin (5-HT) secretion. **a**, Schematic showing fast cyclic voltammetry (FSCV) measuring real-time 5HT secretion from CHGA-mNeon fluorescent cells in human colonoids, where extracellular 5-HT is detected as a change in current at a carbon fibre microelectrode (CFM). **b**, Representative current-time plots of each organoid, after treatment of 10 µM FSK/IBMX, 100 nM Phoenixin-20, or 100 ng/mL IL-13. **c**, Quantified frequencies of 5-HT release events. **d,** The peak amplitude indicating the amount of 5-HT released per event.

To evaluate the functional significance of GPR173, which we found to be exclusively expressed by ECs in mouse and humans, we used its recently identified ligand PNX-20 and FSCV. The frequency of serotonin release events increased following PNX-20 treatment (**Fig. 6c**). However, the peak amplitude, corresponding to the amount of 5-HT released, was not affected (**Fig. 6d**). These results suggest that PNX increases the frequency of 5-HT release events but does not affect the amount of 5-HT secreted during each event.

## Discussion

EECs are a potentially rich source of novel drug targets for the treatment of numerous diseases. However, human EEC biology and their therapeutic control have remained challenging to characterise beyond the well-studied incretin axes. By leveraging advanced methodologies, including CRISPR-Cas9 gene editing and organoid models, we and others have begun to address longstanding challenges in accessing and studying human EECs at high resolution ^67,68,69^. By generating human CHGA-mNeon colonoids and developing a small molecule EEC differentiation protocol, we facilitated the isolation and visualization of EEC populations for transcriptomic profiling. In addition, we utilized TPH1-CFP and PYY-GFP transgenic mouse models to enable comparative analyses of EEC subtypes across species. This dual-species approach allowed us to map both conserved and species-specific features of EEC biology. By integrating bulk and single-cell RNA sequencing with functional assays we sought to delineate the molecular landscape of colonic EECs. This approach led to the identification of a broad array of previously unreported neuropeptides, hormones, and receptors in human colonic EECs. These findings expand our understanding of the signalling pathways that govern EEC function and highlight novel candidates for further exploration. Moreover, we corroborate the presence of key chemosensory receptors, such as the bile acid receptor GPBAR1 and free fatty acid receptor FFAR2, reaffirming the critical role of EECs in nutrient sensing and gut physiology.

In addition to their roles in nutrient sensing, hormone secretion, and gut motility, EECs are increasingly recognized for mediating immune-gut communication. Though their immunological functions remain relatively underexplored, we show that human colonic EECs express receptors and signalling molecules associated with immune function, including cytokine and chemokine receptors. This suggests that EECs sense inflammatory signals and modulate immune responses via hormone and peptide release. For example, we demonstrate that IL-13 enhances GLP-1 and serotonin secretion, indicating that EECs may link immune regulation, particularly TH2 responses, with metabolic processes and or gut defence. Further research is needed to clarify this crosstalk and its role in disease.

We employed fast-scan cyclic voltammetry (FSCV) for the first time in gut organoids to validate serotonin secretion from ECs. FSCV provides unparalleled temporal resolution, allowing for the detection of rapid and transient changes in serotonin release that are often undetectable with conventional methods such as ELISA. Using this approach will facilitate a deeper understanding of the finely tuned regulatory mechanisms governing serotonin secretion from ECs. For instance, we identified GPR173 as a novel receptor exclusively expressed by ECs. FSCV enabled us to demonstrate that the GPR173 ligand, Phoenixin-20, modulated serotonin release by increasing the frequency of release events. These findings suggest that GPR173 may influence gut motility, inflammation, or mood regulation, given the established role of serotonin in the gut-brain axis.

Our interspecies transcriptomic comparisons revealed both conserved and divergent expression patterns, underscoring the limitations of exclusively using animal models for studying EEC biology. Conserved expression was observed for receptors such as NPY1R and FFAR2 whereas receptors like GPR3, GPR160, ULBP1, RXFP4 displayed human-specific expression, highlighting species-specific differences. This emphasizes the importance of conducting direct studies on human tissues to gain a comprehensive understanding of EEC function, particularly in the context of drug development. Additionally, the cross-species catalogue enables novel targets to be identified which can be further explored in mouse models of disease.

Overall, we provide a transcriptomic catalogue of human colonic EECs and the functional validation of two novel receptors involved in EEC biology. The identification of previously uncharacterized receptors, channels and transporters in the database presents the field with an opportunity to identify novel therapeutic interventions in metabolic, gastrointestinal, and inflammatory diseases. Future research should prioritize *in vivo* validation of targets within disease-specific contexts where gut hormones play key regulatory roles. Finally, further investigation of the roles of IL-13Rα1 and GPR173 will lead to a deeper understanding of how these receptors influence gut homeostasis and their therapeutic potential in various pathologies.

## Methods

### Isolation and culture of human colonoids

The isolation of human colonoids was adapted from previously described methods ^14,70^. Three human colonoid lines were derived from colonoscopy biopsies of a 50-year-old male patient (unique identifier FG), a 75-year-old female patient (unique identifier MW) and 33-year-old female patient (unique identifier ZB), respectively, from Denmark Hill NHS Foundation Trust.

The biopsies were rinsed with ice-cold D-PBS (D8537, Sigma-Aldrich) to remove any blood or debris and then incubated in 5 mL 10 mM 1,4-dithiothreitol (DTT; 10197777001, Sigma-Aldrich) for 5 min at room temperature, repeated once with fresh DTT. The biopsies were then incubated in 8 mM EDTA (15575-038, Invitrogen) in D-PBS on a rotator at 4 °C for 1 hr, and subsequently vigorously shaken in cold D-PBS to release crypts. The supernatant containing crypts were collected and centrifuged at 400 rcf for 3 min and were washed in cold D-PBS for three times. 200 crypts per 25 µL were seeded in Cultrex Basement Membrane Extract (BME) (3536-005-02, Bio-Techne) in 48-well plate (Nunc). BME was polymerized for 15 min at 37 °C, then overlaid with 250 µL/well IntestiCult™ Organoid Growth Medium (06010, STEMCELL) supplemented with 100 units/mL Pen-Strep and 10 µM Y-27632 (Y0503, Sigma-Aldrich). Plated organoids were maintained in a 37°C incubator with 5% CO_2_, and the media were changed every other day.

When the crypts grew into organoids, they were maintained in the stem cell medium IFE. IFE medium consists of Advanced DMEM/F-12 (12634, ThermoFisher Scientific), 2 mM GlutaMAX (35050061, Gibco), 10 mM HEPES (15630056, Gibco), 100 units/mL Pen-Strep, 1× B27 supplement (17504044, Gibco), 1× N2 supplement (17502048, Gibco), 0.15 nM Wnt Surrogate-Fc Fusion Protein (N001, ImmunoPrecise Antibodies), 10% R-spondin-1 conditioned medium (in house), 1% Noggin-Fc fusion protein conditioned medium (N002, ImmunoPrecise Antibodies), 50 ng/mL recombinant human EGF (AF-100-15, PeproTech), 1.25 mM N-Acetylcysteine (A9165, Sigma-Aldrich), 10 nM Gastrin (G9145, Sigma-Aldrich), 500 nM A83-01 (2939, Bio-techne), 100 ng/mL recombinant human IGF-1 protein (590904, Biolegend) and 50 ng/mL recombinant human FGF-2 protein (100-18B, PeproTech). Maintained in IFE medium, the human colonoids were passaged every 7 days by mechanical dissociation, at a 1:6 split ratio.

The differentiation of human colonoids was initiated on day 4 after splitting, when IFE medium was substituted by IF* medium till day 11. In the IF* medium, the Wnt Surrogate protein was reduced to 0.045 nM and EGF was removed to accelerate enteroendocrine differentiation. A 48-hr pulse treatment with a combination of small molecules was applied for enhanced differentiation. This treatment involved the use of 10 μM iNotch DAPT (D5942, Sigma-Aldrich), 500 nM iMEK PD0325901 (Sigma-Aldrich) and 40 µM ISX-9 (4439/10, Bio-Techne), which were applied 4 days before the end of the culture process ^14^. After 48 hrs, the treatment was discontinued, and the medium was changed to IF* medium for an additional 48 hrs.

### Transgenic mice

The transgenic PYY-GFP mice (a kind gift from Prof. Rodger Liddle, Duke University) ^63^ were maintained under regulated temperature (21-23°C) and light (12:12 hr light/dark cycle) conditions, with access to *ad libitum* water and chow diet. These mice were housed in compliance with Home Office UK regulations.

### Culture of mouse colonoids

The transgenic Tph1-P2A-iCreERT2 mouse colonoids are a kind gift from Dr Cordelia Imig, University of Copenhagen ^44^. Mouse colonoids were isolated from 8- to 12-week-old male mice and cultured as described previously ^70,71^. The WENR medium consists of Advanced DMEM/F-12, 2 mM GlutaMAX, 10 mM HEPES, 100 units/mL Pen-Strep, 1× B27 supplement, 1× N2 supplement, 50 ng/mL recombinant human EGF, 1.25 mM N-Acetylcysteine, 1% Noggin-Fc fusion protein conditioned medium (N002, ImmunoPrecise Antibodies), 0.15nM Wnt Surrogate-Fc Fusion Protein (N001, ImmunoPrecise Antibodies) and 10% R-spondin-1 CM (in house). Maintained in WENR medium, the colonoids were passaged every 7 days by mechanical dissociation at a 1:6 split ratio.

The differentiation of mouse colonoids was initiated on day 3 after splitting, when WENR medium was substituted by ENR medium till day 7. The ENR medium has the same composition as WENR, but without Wnt Surrogate. A 48-hr pulse treatment of 40 µM ISX-9 was applied from day 3 to day 5 in ENR. After 48 hrs, the treatment was discontinued, and the medium was changed to ENR medium for an additional 48 hrs.

### Generation of human CHGA-mNeon colonoids

The human colonoids (unique identifier FG) were used to generate CHGA-mNeon reporter organoids by CRISPR-HOT. The three plasmids comprising the CRISPR-HOT system for targeting the endogenous CHGA gene locus include the target selector pSPgRNA (Addgene #47108), with sgRNA (CAGCTGCAGGCACTACGGCGGGG) inserted using a previously described protocol ^72^, the frame selector pCas9-mCherry-Frame +1 (Addgene plasmid #66940), and the universal donor pCRISPaint-mNeon (Addgene #174092) ^9^. These three plasmids were kindly provided by Prof Hans Clevers’ group ^8^. Additionally, an EF1α promoter-guided puromycin resistance gene cassette was amplified from the PB513B-1 plasmid (System biosciences) and cloned into the pCRISPaint-mNeon plasmid for antibiotic selection. Briefly, the pCRISPaint-mNeon plasmid was linearized by Q5® Hot Start High-Fidelity 2X Master Mix (M0494S, NEB). The EF1α promoter and the puromycin resistance gene fragments were amplified by touchdown PCR ^73^ using Q5® High-Fidelity DNA enzyme (M0491, New England BioLabs), each with a 20 bp extension complementary to the ends of the linearized pCRISPaint-mNeon plasmid. NEBuilder® HiFi DNA Assembly Master Mix (E2621S, New England BioLabs) was then used for ligation of the two DNA fragments. The ligated fragment was cloned into pCRISPaint-mNeon plasmid using the In-Fusion® HD Enzyme Premix (011614, Clontech). The generated plasmid was named pCRISPaint-mNeon-Efα-PuroR. PCR primers used in amplification and cloning are provided (**Table. S5**).

The three plasmids were transfected to human colonoids by electroporation, adapted from previously reported protocols ^74,75^. Briefly, before electroporation, human colonoids were cultured in IFE medium supplemented with 10 µM Y-27632 and 5 µM Prostaglandin E2 (PGE2) (CAY14010, Cayman Chemical) ^76^ for 3-5 days after splitting. Ten wells of organoids were collected and mechanically broken by vigorous trituration with a p-200 micropipette. The pelleted organoids were then resuspended in 1 mL TrypLE Express Enzyme with 10 µM Y-27632, and incubated in 37 °C water bath for 3 min to dissociate into clusters of 10-15 cells ^75^. The dissociated organoids were washed in Opti-MEM (31985062, Gibco) twice. Every 1×10^5^ to 5×10^5^ cells were resuspended in 100 µL BTXpress buffer, containing 5 µg of each plasmid, 15µg in total. After that, the cell-plasmid mixture was placed into a 2-mm electroporation cuvette, and the electroporation performed immediately before the cells precipitated, using the NEPA21 electroporator (Nepa Gene). After electroporation, the cells were seeded in IFE medium with 10 µM Y-27632 and 5 µM PGE2. Five days after electroporation, cells were selected with 1 μg/mL puromycin in IFE medium.

### Whole mount immunostaining of colonoids

Human colonoids embedded in BME were fixed in 4% formaldehyde (28908, Thermo Fisher) for 45 minutes at room temperature. After fixation, the organoids were washed in ice-cold D-PBS containing 2% BSA (A7906, Sigma-Aldrich). The organoids were incubated on a rotator at room temperature for 1 hr in blocking/permeabilization buffer, which consisted of 2% BSA, 5% donkey serum (D9663, Sigma-Aldrich) and 0.5% Triton X-100 (X100, Sigma) in D-PBS. After the blocking/permeabilization step, the organoids were incubated overnight at 4 °C in a rotator with primary antibodies rabbit polyclonal anti-CHGA (1:800; ab15160, Abcam). On the next day, the organoids were washed and incubated with secondary antibodies, Alexa Fluor™ 568 donkey anti-rabbit (1:500, Invitrogen) in a rotator for 1 hr at room temperature. Nuclear counterstaining dye, Hoechst 33342 (5 µg/mL; H3570, Invitrogen) was added to the secondary antibody solution and incubated for another 15 min. Following thorough washes in D-PBS to remove all unbound antibodies, the organoids were mounted on a glass slide with a drop of mounting medium Fluoromount-G^®^ (0100-01, Cambridge Bioscience) and covered with a coverslip. The slides were air-dried in the dark overnight at room temperature before being analysed by confocal microscopy.

### Immunohistochemical staining of colon biopsies

Paraffin-embedded tissue blocks of human colon mucosal biopsies from healthy patients were provided by Dr Polychronis Pavlidis (King’s College Hospital, Denmark Hill). Tissue sections (5 μm) were mounted on positively coated Superfrost® Plus slides (MIC3040D2, Scientific Laboratory Supplies). Sections were deparaffinized and subsequently underwent heat-induced antigen retrieval by microwaving in 10 mM sodium citrate buffer (pH 6.0) for 10 min. After rinsing with water, slides were incubated at room temperature for 30 min in a blocking/permeabilization buffer containing 2% BSA, 5% donkey serum, and 0.5% Triton X-100 in D-PBS. Following blocking, sections were incubated overnight at 4 °C in a humidified chamber with primary antibody IL-13Rα1 (1:200, rabbit; ab79277, Abcam) and GLP-1 (1:200, mouse; ab23468, Abcam) or 5-HT (1:200, goat; 20079, Immunostar). Slides were then washed in D-PBS and incubated with secondary antibodies (1:500, Alexa Fluor™, Invitrogen) for 1 hr at room temperature. Both primary and secondary antibodies were diluted in blocking/permeabilization buffer. After the secondary antibody incubation, slides were washed with D-PBS and mounted with DAPI Fluoromount-G^®^ (0100-20, Cambridge Bioscience). The slides were air-dried in the dark overnight at room temperature before being analysed by confocal microscopy.

### FACS sorting fluorescent EECs

After differentiation of the colonoids, they were dissociated to single cells in 2-4 mL TrypLE Express (Gibco), with 4 µg/mL DNase (79254, Qiagen) at 37 °C for 15-20 min. Dissociated single cells were then filtered through a 40 µm cell strainer. The cells were then resuspended in FACS buffer (Advanced DMEM/F-12 medium, 2 mM GlutaMAX, 10 mM HEPES, 10 µM Y-27632, 2% FBS and 2 mM EDTA). For CHGA-mNeon and PYY-GFP cells, 0.1 µg/mL DAPI (10236276001, Merck) was used as the live/dead cell marker. For TPH1-CFP cells, 7-AAD (00-6993-50, Invitrogen™) at 0.25 µg per million cells was used. FACS was performed on the BD FACS Aria™ II (Beckton Dickinson) with a nozzle size of 100 µm. The gating strategy images were generated by FlowJo software.

### RNA extraction and real-time quantitative PCR (RT-qPCR)

FACS-sorted cells were collected in a LoBind tube containing 500 μL of D-PBS. The cells were then centrifuged at 600 rcf, 4 °C for 5 min. The total RNA extraction of the cells and cDNA transcription were performed using the SYBR™ Green Fast Advanced Cells-to-CT™ Kit (A35379, Invitrogen). RT-qPCR was performed using PowerUp SYBR Green Master Mix (A25742, Thermo Fisher) with QuantiTect primers or 500nM designed primers (**Table. S3**) on LightCycler 96 (Roche).

Total RNA extraction from the whole organoids was conducted using the RNeasy Kit (74106, Qiagen) after they were released from BME using Cell Recovery solution (11543560, Corning). On-column DNase I (79254, Qiagen) was used to remove residual genomic DNA. cDNA transcription was performed using Power High-Capacity cDNA Reverse Transcription Kit (4368813, Applied Biosystems). RT-qPCR was performed with QuantiFast SYBR Green PCR Kit (204057, Qiagen) with QuantiTect primers or 500nM designed primers (**Table. S3**) on a LightCycler 96 (Roche).

The relative gene expression levels were determined by averaging the Ct values from technical duplicates for each biological sample and then normalizing them against the expression of the reference gene beta-2-microglobulin (*B2M*).

### Single cell RNA-sequencing and data analysis

Fluorescent single cells were sorted into 384-well cell capture plates (HSP3801, Bio-Rad) for SORT-seq. The samples were processed by Single Cell Discoveries (Utrecht, Netherlands) for library preparation, sequencing and alignment, following the published SORT-seq protocol ^35^. The scRNA-seq analysis of raw counts was conducted in RStudio using the R package Seurat v4.3.0 ^36^.

For human CHGA-mNeon cells, those with fewer than 1550 transcripts, unique feature counts exceeding 2500 or falling below 200, or mitochondrial counts exceeding 30% were excluded. The cells were normalized using SCTransform ^77^. After normalization, distinct cell clusters were identified through shared nearest neighbour clustering optimization, based on the 4 most variable dimensions. A resolution of 0.4 was used during the clustering process. For mouse TPH1-CFP cells, those exceeding 4000 unique feature counts or falling below 200, or with mitochondrial counts surpassing 30%, were excluded. Following normalization using SCTransform, shared nearest neighbour clustering optimization was applied based on the 5 most variable dimensions, with a resolution of 0.4. This process identified three distinct cell clusters. However, one cluster was enriched with genes associated with cell death, indicating cell damage. Therefore, this population was removed. For mouse PYY-GFP cells, those with over 2500 unique feature counts or less than 200, along with mitochondrial counts exceeding 5%, were excluded. After SCTransform normalization, shared nearest neighbour clustering optimization with the 4 most variable dimensions and a resolution of 0.5 identified two distinct cell clusters.

The cell clusters were plotted using UMAP dimension reduction. Expression of genes of interest across different cell clusters were plotted on heatmaps or distribution plots. Gene expression table of distinct clusters were generated by AverageExpression. Co-expression of two features was visualized by FeaturePlot.

### Bulk RNA-sequencing and data analysis

Around 100,000 fluorescent and non-fluorescent single cells were FACS-sorted, pelleted and underwent RNA extraction with TRIzol™ LS Reagent (10296010, Invitrogen™). Three replicate samples were prepared for both fluorescent and non-fluorescent groups. The RNA samples were processed by Single Cell Discoveries for cDNA library preparation and sequencing. 75-bp paired-end sequencing of the cDNA libraries was performed on an Illumina NextSeq™ 500 platform. After sequencing, the paired-end reads were aligned to the human reference genome GRCh38/hg38 or mouse reference genome GRCm38/mm10 using Burrows-Wheeler Aligner (BWA) ^78^.

Differential gene expression analysis between the fluorescent and non-fluorescent groups (n=3) was performed in RStudio using the Bioconductor package DESeq2 v1.38.1 ^79^. The PCA plot of differentially expressed genes was generated to demonstrate clusters of samples based on their similarity. Gene lists encoding receptors, orphan GPCRs, transporters, ion channels, gut hormones, transcription factors, cytokines were generated from the highly expressed gene list using QIAGEN Ingenuity Pathway Analysis (IPA) software, with log2FC > 1.5 and p-value < 0.05. Specifically, the genes with predicted protein location in the extracellular space were included in the gut hormone gene list. For display in heatmaps, genes were ranked by fold change compared against non-fluorescent cells. Differentially expressed genes with a p-value of 0 were replaced with 3×10^-300^ to enable visualisation in the volcano plot.

### Measurement of GLP-1 secretion in human colonoids

Differentiated human colonoids were harvested with cell recovery solution and washed with D-PBS, with one well as one sample. The organoids were then incubated for 2 hrs at 37 °C in 100 µL of saline secretion buffer containing 138 mM NaCl, 4.5 mM KCl, 4.2 mM NaHCO_3_, 1.2 mM NaH_2_PO_4_, 2.6 mM CaCl_2_, 1.2 mM MgCl_2_, 10 mM HEPES (pH 7.4), with freshly added 0.1% fatty acid free BSA (A6003, Sigma-Aldrich) and 50 µM DPP IV Inhibitor (DPP4, Sigma-Aldrich). Four groups were set up: a negative control, treatment of 10 µM Forskolin (FSK, F6886, Scientific Laboratory Supplies) and 10 μM IBMX (I5879, Merck), and treatments with 20ng/mL or 100 ng/mL recombinant human IL-13 protein (R&D Systems, 213-ILB-010). The supernatant and organoid lysates were collected to measure GLP-1 concentrations in secretion and total protein concentrations, respectively. Organoid lysates were prepared in 100 μL of D-PBS with protease inhibitors (A32955, Thermo Fisher) and sonicated on ice for 30 seconds at an amplitude of 10-14. The homogenates were then centrifuged at 5,000 rcf, 4 °C for 5 min, and the supernatant was collected as organoid lysate samples. Both secretion and lysate samples were stored at −70 °C. The GLP-1 concentrations of secretion and lysate samples were measured using GLP-1 ELISA kit (Merck, EGLP-35K). Total protein concentrations of organoid lysates were measured using Pierce™ BCA Protein Assay kit (23227, Thermo Fisher). The relative GLP-1 secretion levels were determined by averaging the GLP-1 concentrations from technical duplicates for each biological sample and then normalizing them against the concentrations of the total protein.

### FSCV cell measurements of 5-HT secretion in human colonoids, data acquisition and analysis

CFM fabrication and FSCV data acquisition was performed as described previously ^10^. Briefly, carbon fibre (T-650, 7 µm diameter; Goodfellow) was aspirated into a glass capillary (1.0 mm outer diameter, 0.58 mm inner diameter) and the capillary was pulled using PE-22 micropipette puller (Narishige Group) to create a seal around the fibre. The exposed carbon fibre was manually trimmed to 100 µm (+/− 2 µm) in length and back-connected using a pinned stainless-steel wire. The carbon surface was coated with Nafion™ (Liquion Solution, LQ-1105, 5% by weight Nafion, Ion Power) by applying 1 V (vs. Ag/AgCl) for 30 sec. All FSCV measurements were acquired in the saline secretion buffer as described above. Data collection and analysis was performed with WCCV 3.06 software (Knowmad Technologies), Dagan potentiostat (Dagan Corporation), and Pine Research headstage (Pine Research Instrumentation). A waveform optimized for 5-HT detection was applied (0.2 V to 1.0 V to −0.1 V to 0.2 V vs. Ag/AgCl at 1000 V/s) and measurements were taken at 10 Hz for 30 sec per file.

For FSCV measurement, differentiated human CHGA-mNeon colonoids embedded in BME were mechanically detached from the plate using a p-1000 micropipette, washed in D-PBS to remove all culture medium, and then resuspended in 2 mL of saline secretion buffer. They were transferred to a 3.5 cm-dish, which was then affixed to the Bio Station IM (Nikon). EC cell-enriched structures were identified by mNeon green fluorescence before shielding the setup with aluminium foil grounded as a Faraday cage. The Ag/AgCl-reference electrode was placed in the dish and the CFM was positioned using a QUAD micromanipulator (Sutter Instruments). Repeated 30 sec-files were taken to record spontaneous activity for a total of 15 min. Subsequently, 100 nM PNX-20 or 100 ng/mL IL-13 was added, and recordings continued immediately for an additional 15 min. Events releasing 5-HT were identified by the 5-HT-specific oxidation peak in the cyclic voltammogram and quantified for each file. Frequency of 5-HT release event was calculated for each 30 sec-file, and peak amplitude was determined as the current difference immediately before release and the peak.

### Proteomics mass spectrometry measurement

Around 100,000 sorted CHGA-mNeon^+ve^ and mNeon^-ve^ cells were pelleted and lysed with Branson sonifier 150 in 8.0M urea buffer. 20 µg of proteins from each sample were reduced with 7mM of 1,4-dithiothreitol for 1 hour at 56 °C and alkylated with 12.5 mM iodoacetamide for 1 hour at room temperature in the dark. Samples were digested with trypsin enzyme at 1:50 ratio (enzyme:protein) and incubated at 37 °C overnight. The digestion was quenched with 1% formic acid and tryptic peptides were purified and desalted by C18 spin column. Peptides were dried to completion by SpeedVac (Thermo) and resuspended in a loading buffer (2% acetonitrile in 0.1% formic acid) for LCMS analysis.

The tryptic peptides were subjected to LCMS system for analysis. For liquid chromatography, a reverse phase Thermo Acclaim Pepmap trap column (2 cm length, 75 µm in diameter and 3 µm C18 beads) were connected to a nanoflow HPLC (RSLC Ultimate 3000) on an Easy-spray C18 nano column (50 cm length, 75 µm in diameter, ThermoFisherScientific). Buffer A (5% ACN, 0.1% formic acid) and buffer B (80% ACN, 0.1% formic acid) were used. Peptides were eluted with a linear gradient of 5%–55% buffer B at a flow rate of 250 nl/min over 100 min at 45 °C. Peptides were directly ionized within the easyspray ion source (Thermo) and injected into Orbitrap Fusion Lumos mass spectrometers (Thermo).

MS data generated were collected within Xcalibur 4.4 to acquire MS data using a “Universal” method by defining a 3s cycle time between a full MS scan and MS/MS fragmentation. We acquired one full-scan MS spectrum at a resolution of 120,000 at 200 m/z with a normalized automatic gain control (AGC) target (%) of 250 and a scan range of 300∼1600 m/z. The MS/MS fragmentation was conducted using CID collision energy (35%) with an orbitrap resolution of 30000 at 200 m/z. The AGC target (%) was set up as 200 with a max injection time of 128ms. A dynamic exclusion of 30s and 2-7 included charged states were defined within this method.

Resulting raw files were searched again the human Uniprot fasta database within Thermo Proteome Discoverer (PD, version 2.5) allowing two missed trypsin cleavage sites and methionine oxidation. Carbamidomethylation on cysteine residues was set as a fixed modification. Precursor mass tolerance was set as 20 ppm and fragment ion tolerance was set as 0.6 Da. Peptides that met the false discovery rate cut-off of 1% based on the searching again a decoy database was considered for further analysis. Precursor ion intensities extracted from PD were applied for differential analysis using two-sample t-test algorithm embedded in Perseus software v2.1.1.0 using two-sample t-test algorithm ^80^. Accession IDs were converted to gene IDs by SynGO ^81^.

### Statistics

The unpaired t-test was utilized to compare two unrelated groups when the test statistic followed a normal distribution. For comparisons involving multiple conditions, the one-way analysis of variance (ANOVA) was conducted.

Statistical significance was accepted at p values < 0.05. The data were presented as mean ± SEM (n ≥ 3) and denoted as *p < 0.05, **p < 0.01, ***p < 0.001, or ****p < 0.0001 to indicate the level of significance. GraphPad Prism 10 was used for generating graphs and performing statistical analysis.

---

**Fig. S1 a,** Principal component analysis (PCA) plot comparing mNeon-positive and -negative cell populations by LCMS intracellular proteomics analysis (n=4). **b,** Volcano plot showing differential expression of selected proteins in CHGA-mNeon cells. **c,** Heatmaps showing different categories of genes enriched in human colonic EECs with the highest fold change and p values < 0.01, including cell surface proteins (orphan GPCRs, transporters, ion channels), and gut hormones, transcription factors, cytokines.

**Fig. S2 a,** Heatmaps showing different categories of genes enriched in mouse colonic ECs with the highest fold change and p values < 0.01, including cell surface proteins (orphan GPCRs, transporters, ion channels), and transcription factors, cytokines. **b,** Venn diagrams showing the conserved and uniquely expressed genes in different categories between the human colonic EECs (blue) and mouse colonic EC cells (green) based on bulk RNA sequencing analysis.

## Author contributions

Y.L. and G.A.B. conceptualized the project and wrote the manuscript. Y.L. generated the reporter organoid model and analysed the EEC transcriptome database. B.B. and P.H. performed the FSCV serotonin secretion experiments and analysis. L.M. and K.G.M. participated in the editing of the manuscript. M.J. participated in bulk RNA analysis. N.H. participated in plasmid cloning and confocal imaging. X.Y. conducted the proteomics study. B.H. provided human biopsies for isolation of organoids. All authors reviewed the manuscript.

## Acknowledgements

We would like to thank Dr Joep Beumer and Dr Hans Clevers from Hubrecht Institute for providing the CRISPR-HOT plasmids, Dr Cordelia Imig from University of Copenhagen for providing TPH1-CFP mouse colonoids and Dr Polychronis Pavlidis from King’s College Hospital for providing paraffin-embedded tissue blocks of human colon mucosal biopsies. We appreciate the assistance from the staff in BRC Flow cytometry core, Guy’s and St Thomas NHS Foundation Trust, Nikon Imaging Centre and Proteomics Facility at King’s College London. Y.L. acknowledges support from China Scholarship Council (K-CSC) and the Society for Endocrinology (Early Career Grant). K.G.M is supported by Diabetes UK (18/0005886, 20/0006295), the BBSRC (BB/W001497/1, BB/X017273/1). Both G.A.B and K.G.M are supported by the MRC (MR/Y013980/1) and the Wellcome Trust (310835/Z/24/Z).

