## Supplementary figures and images for "Mapping the druggable targets displayed by human colonic enteroendocrine cells"

### Supplementary Figures

#### Supplementary Fig 1

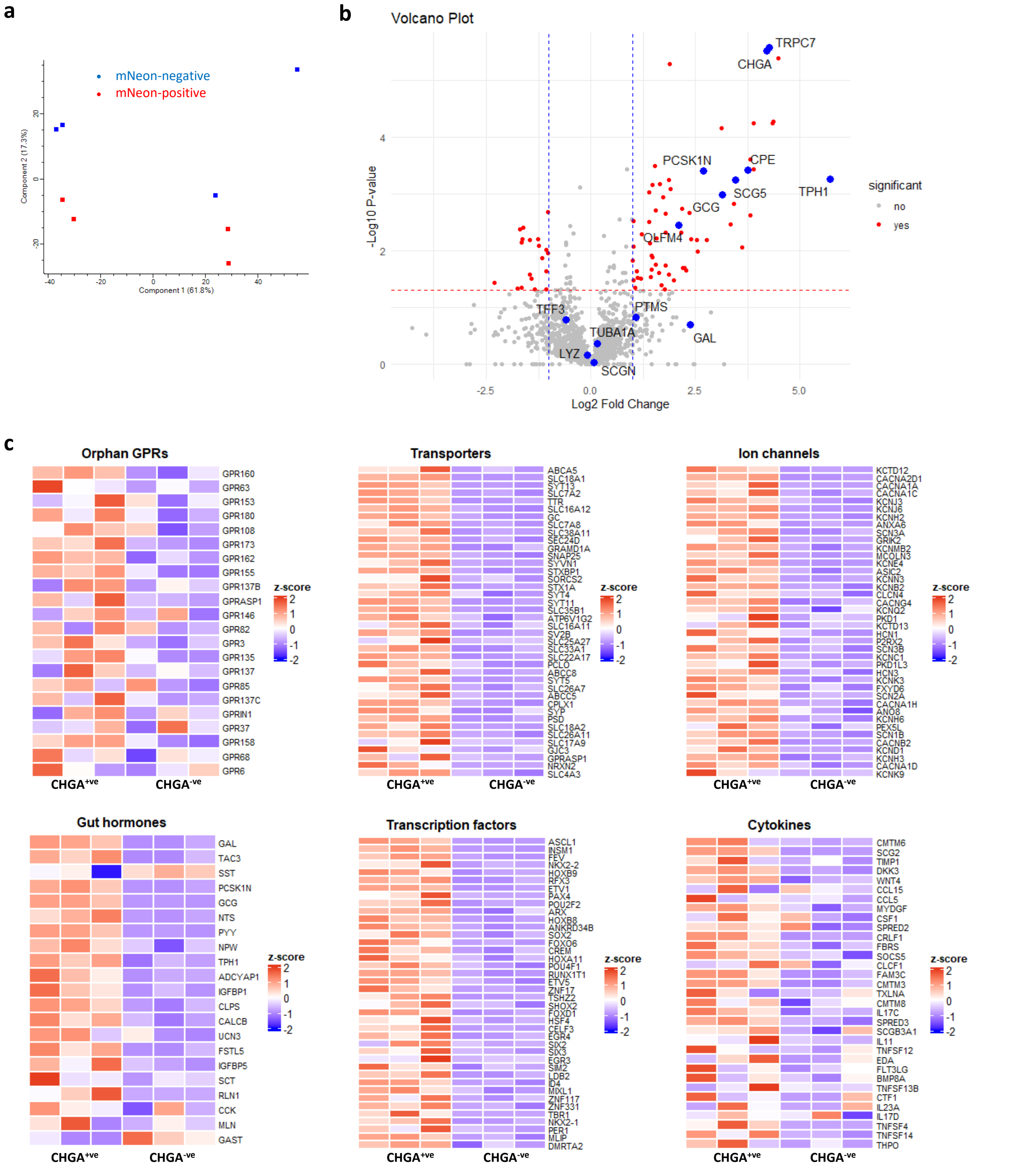

#### Supplementary Fig 2

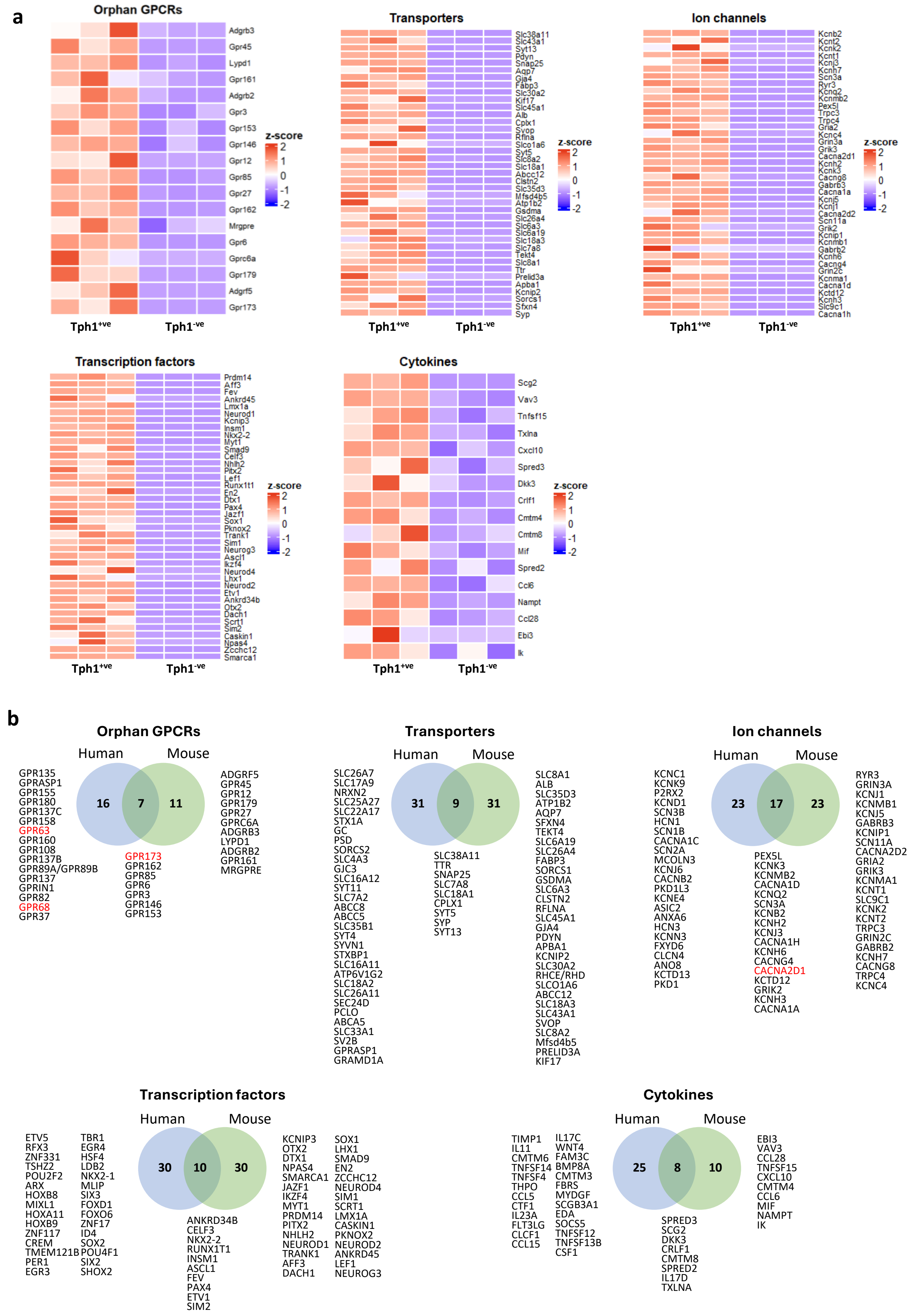
